# Temporal rules of fear memory cooperation and competition

**DOI:** 10.1101/2024.05.28.596109

**Authors:** Natália Madeira, Inês Campelo, Rosalina Fonseca

**Affiliations:** Learning and Plasticity, i3s – Instituto de Investigação e Inovação em Saúde, Universidade do Porto; Porto, Portugal; Cellular and Systems Neurobiology, NOVA Medical School, Universidade NOVA de Lisboa; Lisboa, Portugal

## Abstract

Memory consolidation is highly influenced by ongoing experiences. We explore the temporal rules that determine whether events are cooperatively associated or competitively separated. We show that neutral events are associated with fearful events if they occur within less than 30 minutes. In some individuals, memory association can lead to a competitive suppression of the fearful response by the neutral event. Activation of the thalamic MGm inputs to the lateral amygdala, led to an increase in memory association, whereas manipulation of the cortical inputs had no effect. Introducing a third event leads to competition depending on the temporal relationship between the initial association and the competitive event. Our results show a critical temporal rule of memory association, modulated by thalamic activity that shapes fear memory consolidation.

## Introduction

The ability to acquire and retrieve memories is fundamental to adequate behavior to environmental challenges. It is now well accepted that the acquisition of long-term memories (LTM) involves a process of consolidation, in which labile memories, sensitive to disruption, become stabilized as long-lasting traces^1^. However, memories are not immutable traces but rather constantly changing representations^2^. Indeed, remembering a previously acquired memory reactivates the trace, returning memories to this active state, during which previously acquired memories can be modified by temporally occurring events. In other words, even if each learning event can be seen as a “new” memory formation process, considerable evidence indicates that new memories are formed in an interleaved fashion, upon a large network of pre-existing knowledge. This implies that learning is highly influenced by previous experience and that previously consolidated memories can be reactivated during new learning^3,4^. Thus, as long as memories are in the active state, either during acquisition or after reactivation, their content can be modified by concurrent events. Consistent with this, previous studies, have shown that at the time of acquisition, two temporally separated events can be linked and remembered as a single memory trace^5,6^. Additionally, exposing animals to a novel environment, before or after a learning event acts as a positive reinforcer of memory consolidation^7^. These results show that distinct events, including an unexpected novel event, can be associated in time promoting a cooperative memory consolidation. As important as associating events in time, the ability to distinguish events is highly relevant. Even more if one considers aversive events and indeed the inability to distinguish between aversive versus neutral events leads to a generalized anxious response that is maladaptive^8^.

In auditory fear-conditioning, subjects learn to associate a neutral stimulus (conditioned stimulus - CS) with a coincident aversive stimulus (unconditioned stimulus - US^9,10^). Once conditioned, the neutral stimulus will evoke a conditioned response (CR) resulting in immobilization or freezing in rodents, providing a behavioral measure of the learned association. Conversely, in auditory discriminative learning, the animal learns to distinguish between two neutral stimuli, a CS+ paired with the US and a CS-unpaired. As the animal learns this task, responses of amygdala pyramidal neurons increase upon CS+ presentation and decrease upon CS-presentation^11^. The leading cellular model underlying auditory fear conditioning is a form of Hebbian long-term potentiation (LTP), induced by the association between the auditory thalamic and cortex projections (CS) and the nociceptive input (US)^12^. The auditory information reaches the amygdala via direct projections from the medial division of the medial geniculate nucleus (MGm-thalamic input) and an indirect projection, through the projection of ventral division of the medial geniculate nucleus (MGv – cortical input) to the auditory cortex which in turn, projects to the LA^13,14^. Previous studies have shown that thalamic and cortical input projections contribute differently to the acquisition of auditory discriminative memories. Cortical projections have been associated with the processing of more complex CSs whereas the direct thalamic projections have been considered to be a fast but less accurate information pathway^15^. This is supported by the observation that neurons in the MGv and primary auditory cortex are sharply tuned and tonotopically organized, whereas neurons in the MGm are multisensory and show non-tuned auditory responses. While the activation of either the cortical or thalamic inputs is sufficient for fear-conditioning learning, in auditory discriminative learning, co-activation of both inputs seems to be necessary for discrimination^16^.

As for memory, synaptic inputs can also interact cooperatively and competitively^17–19^. Synaptic cooperation and competition are two forms of heterosynaptic plasticity based on the interaction of biochemical signals that allow synapses to integrate cell-wide activity within large time windows^17,20,21^. Taking advantage of the detailed knowledge of the circuits underlying fear memory acquisition, we have built a detailed model of the temporal rules underlying synaptic cooperation and competition between the cortical and thalamic inputs to the lateral amygdala. We found that cortical synapses can cooperate with thalamic synapses if stimulated within a temporal window of 30 minutes^22,23^. The reverse interaction, in which the thalamic synapses are reinforced by cortical activation is restricted to a much shorter time window of 7.5 minutes due to the activation of cannabinoid receptor type 1 (CB1R). If CB1Rs are inhibited, thalamic synapses can extend the time interval of cooperation to 30 minutes, similar to the time interval observed for cortical synapses. Thus, thalamic-cortical cooperation reinforces both inputs in associative but temporally asymmetrical, manner. This suggests that, once plasticity is induced, ongoing activity and activation of CB1R restricts the time window for thalamic-cortical cooperative plasticity. Thalamic and cortical synapses also engage in synaptic competition^24^. Introducing a third stimulating electrode placed on internal capsule fibers to activate a second thalamic input creates a competitive load by increasing the number of activated synapses that capture PRPs and as for cooperation, time is crucial to induce competition. If the second thalamic weak stimulation was delayed by 30 minutes, no competition was observed, suggesting that synapses are susceptible to perturbation, by competition, for a restricted time window. Inhibition of CB1 receptors promotes competition, possibly by reducing presynaptic glutamate release and synaptic activation^24^. Building on this model of synaptic cooperation and competition between thalamic and cortical inputs to the lateral amygdala, we have designed a modified discriminative learning task that allowed us to test the temporal rules of memory cooperation and competition as well as the contribution of the thalamic and cortical inputs to discriminative fear memory acquisition.

## Results

### Memory cooperation has a time rule

Our synaptic model ^24,25^, predicts that events occurring within a short time window will be associated resulting in a form of memory cooperation, whereas longer time windows will allow event separation. To test this hypothesis, we expose animals to two different events separated by three different time points, 7.5, 30, and 60 minutes. These times were derived from our synaptic analysis of thalamic and cortical inputs converging into the lateral amygdala. Event one consisted of one particular context (A) where an auditory stimulus (CS-) is presented, whereas in event two, occurring in a different context (B), a second auditory stimulus (CS+) is presented co-terminating with a foot shock (Figure 1). As we expected, we found that for the shortest time interval used, 7.5 minutes, the events are associated and animals respond with a freezing response to both stimuli, CS- and CS+ in the test session (Figure 1A). Also, for the longest interval, as we hypothesized, animals can separate the two events and freeze only to the CS+ stimuli (Figure 1C). The cooperative effect seen in the 7.5 minutes interval is not due to an overall generalization of fear, given that, if CS-is not present in the first event, no response is seen in the test trial (Supplementary Figure 1A).

**Figure 1.**
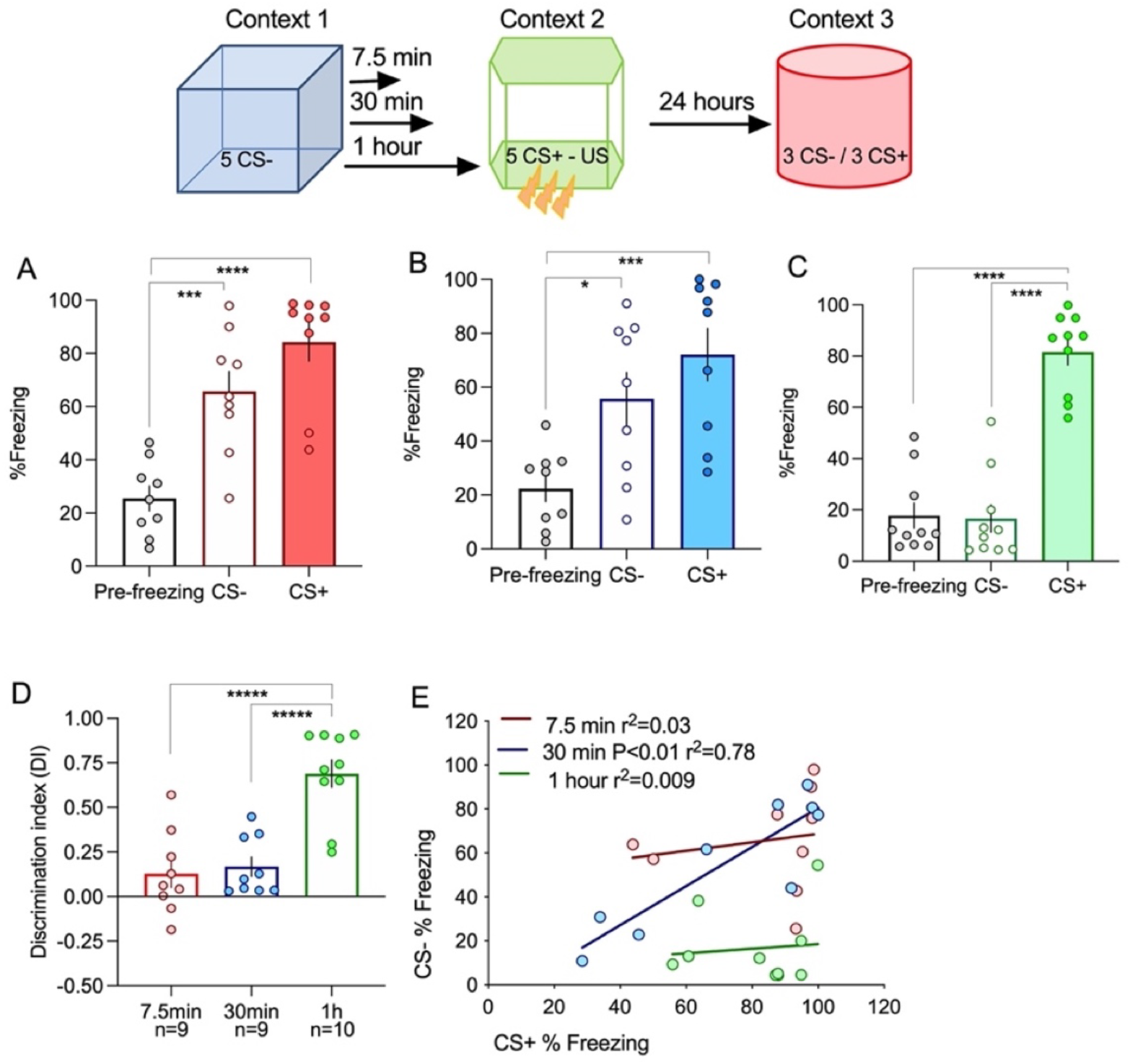
Memory association depends on the time interval between events. Diagram illustrates the sequence of trials during training (day 1) and in test (day 2). A. When events are separated by 7.5 minutes, events are associated and animals significantly freeze above baseline responses to both CSs (ANOVA F=20.87 P<0.0001; ***P=0.0007; ****P<0.00001 n=9). B. Increasing the time interval between event 1 and 2 to 30 minutes also leads to fear memory association (ANOVA F=8.98 P=0.0012; *P=0.026; ***P=0.001 n=9). C. Increasing the time interval to one hour allows the dissociation between event 1 and 2, resulting in a specific fear response to the CS+ (ANOVA F=53.47 P<0.0001; ****P<0.0001; ****P<0.0001 n=10). D. Analysis of the discrimination index shows that only the interval of 1 hour allows the dissociation between the neutral and fearful events (ANOVA F=20.13 P<0.0001; ****P<0.0001; ****P<0.0001). E. Plotting the individual response to the two CSs reveals that for the intermediate time tested (30 minutes) there is a positive correlation between the fear response to the CS- and the CS+, correlation that is not observed in any other time interval tested.

For the 30-minute interval, the average freezing response is similar to the one observed in the 7.5 minutes with significantly higher freezing levels for both CS- and CS+. This is also evident in the analysis of the discriminative index shown in Figure 1D. For the time interval of one hour, the discriminative index is significantly higher than for the other two-time intervals tested. Interestingly, when analyzing individual responses, we found that for the intermediate time interval (30 minutes), animals were distributed in two distinct groups. One group of animals freezes to both stimuli, CS- and CS+, showing memory cooperation, whereas a second group does not freeze to either stimulus. This is better appreciated by plotting the freezing levels to CS+ and CS-of individual subjects, as shown in Figure 1E. For the short (7.5min) and long interval (60min) there is no correlation between CS+ and CS-freezing levels, with values distributing around higher CS-values in the case of the 7.5 minutes (red line) and low CS-values for the 60 minutes interval (green line). In the case of the 30 minutes, we found a significant correlation (blue line) between CS+ and CS-, representing the two groups of responses, higher freezers and lower freezers to both stimuli. This observation suggests that animals in the 30-minute interval generalize their responses, but they can either generalize to freeze or generalize to not freeze.

In our synaptic model, we found that inhibiting endocannabinoid receptors (CB1R) promoted synaptic cooperation by extending the time window in which thalamic synapses cooperate with cortical synapses^25^. To test whether inhibiting cannabinoid receptors also alter our memory cooperation time rule, we inhibited CB1R in the lateral amygdala, during the 60 minutes between events one and two (Supplementary Figure 2A). According to our prediction, we found that inhibiting CB1R increased memory cooperation as seen by an increase in freezing to the CS-(Supplementary Figure 2B/C) and a significant decrease in the discrimination index (Supplementary Figure 2D). The promoting effect of inhibiting CB1R in synaptic cooperation was associated with a modulation of thalamic inputs, by increasing the time windows in which thalamic inputs can cooperate with cortical inputs.

Having this in mind, we used chemogenetic manipulations of the thalamic and cortical inputs to the LA to assess their role in modulating the time rule of cooperation. Transfection of the Mgm thalamic nuclei with a vector expressing the hM3D(Gq) receptor resulted in the labeling of axons targeting the LA region via the internal commissure (Figure 2A/A’). Using the 60-minute interval between events, in which memory cooperation does not occur, we found that injection of CNO before the training trial, increased cooperation resulting in a generalized freezing response to both CS- and CS+ stimuli (Figure 2B-D). Analysis of the discrimination index shows that discrimination is significantly lower if thalamic inputs are activated by CNO injection in the presence of a vector expressing hM3D(Gq), compared to either CNO+open vector or saline+hM3D(Gq) (Figure 2E). Interestingly, when plotting the level of freezing to the CS+ against the freezing response to the CS-, we found that chemogenetic activation of the thalamic inputs results in a significant correlation (Figure 2F), a result similar to the 30-minute interval. This result shows that activating thalamic inputs reduces the time window of memory cooperation as we have observed at the synaptic level. We then asked whether activation of the cortical input also increases memory cooperation. To do this, we transfected the primary auditory region with the hM3D(Gq) receptor-expressing vector (Figure 2G). In this situation, the administration of CNO did not alter cooperation and animals were able to discriminate the two events, showing a preferential freezing response to the CS+ (Figure 2H).

**Figure 2.**
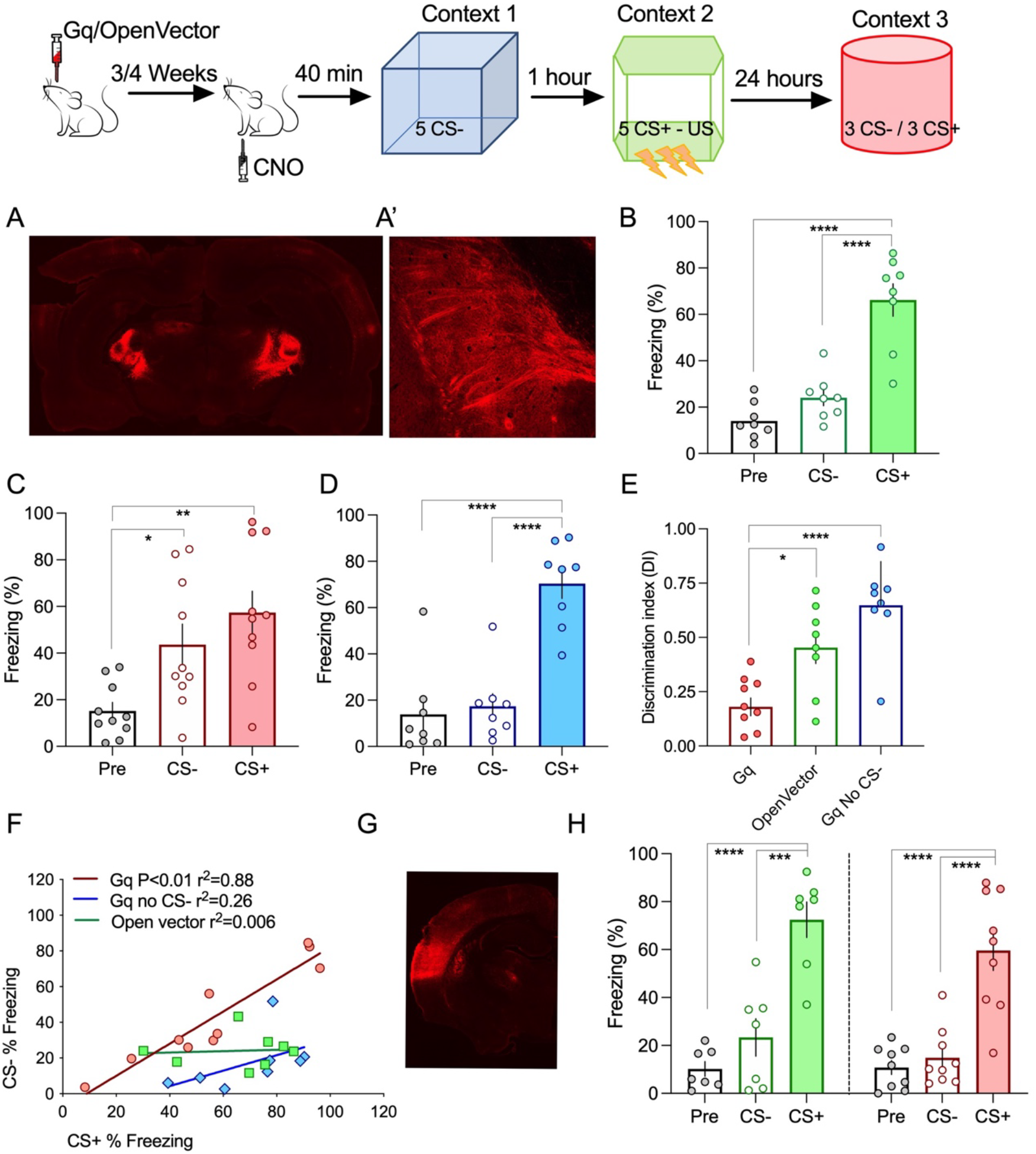
Memory association is influenced by the activity of the thalamic inputs. A/A’. Injection of AAVs in the thalamic medial division of the medial genicular nucleus (MGm and PIN) results in transfection of afferents that project to cortical areas and the lateral amygdala, by the internal capsule. B. Using the one-hour interval, transfecting neurons with an open vector does not alter the ability of animals to distinguish between the neutral and aversive event (ANOVA F=33.73 P<0.0001; ****P<0.0001; ****P<0.0001 n=8). C. Increasing thalamic activity by CNO activation of the Gq DREADD led to memory association even if neutral and aversive event occur with an interval of one hour (ANOVA F=7.88 P=0.002; *P=0.036; **P=0.002 n=10). D. Memory association is not due to the increase of thalamic activation per se since fear responses to the neutral event is dependent on the presenting the CS-during training (ANOVA F=25.74 P<0.0001; ****P<0.0001; ****P<0.0001 n=7). E. Analysis of the discrimination index shows that increasing thalamic activity significantly decreases discrimination (ANOVA F=15.45 P<0.0001*P=0.01; ****P<0.0001). F. Responses to the neutral event (CS-) is correlated to the response of the aversive event (CS+) if thalamic activity is increased. No correlation is observed in animals transfected with the open vector nor in animals where the CS-is not present in training. G. Injection in the primary auditory cortex. H. No differences were observed between animals where the cortical input activity is increased by CNO Gq activation versus animals expressing an open vector (Open Vector ANOVA F=25.83 P<0.0001; ****P<0.0001 ***P=0.0001 n=7; Gq ANOVA F=23.52 P<0.0001; ****P<0.0001 ****P<0.0001 n=9).

### Memory competition is dependent on time and order of events

At the synaptic level, competition was induced by the temporally related stimulation of three inputs to the LA (one cortical, two thalamic)^24^. Given that neurons in the MGm thalamic nuclei have a multimodal response, including visual stimuli, we added a third event in which animals are exposed to a third, different context and visual stimuli (green flashing led). Again, based on our synaptic competition model, we predicted that competition would occur if the third event was temporally close to the two initial events whereas increasing the elapsed time would leave intact the initial memory association. Competition was induced by exposing animals to the light trial 7.5 or 60 minutes after the CS+ and CS-events. In the absence of light, cooperation was observed (Supplementary Figure 3A) whereas that introducing the light at event 3, leads to a mild form of competition, reducing both responses to the CS+ and CS- (Supplementary Figure 3B). As predicted, if the third event occurred 60 minutes after the initial CS+/CS- association, no competition was observed (Supplementary Figure 3C). Given that the initial CS+/CS- association was done with the 30-minute interval, in the absence of the light stimulation we observe that again freezing levels to the CS+ are correlated to CS-freezing responses (Supplementary Figure 3D). Interestingly, although introducing the light resulted in the reduction of freezing responses, the correlation is still present, suggesting that the competitive reduction of freezing responses was done to the CS+/CS- association as a block. In the situation where the light is presented 60 minutes after the initial CS+/CS- association, no change in the correlation was observed. We do not observe any significant freezing response to light in the test trial, suggesting that memory association is not observed between multimodal stimuli in our experimental conditions.

Given that competition by the light interfered with the freezing response of both the CS+ and CS-we designed a second competitive experiment, in which the light event was present in between the two sound events (Figure 3). Again, we used the 30-minute interval between the CS+ and CS-event and the light-event occurred in between so that the time interval between the light-event and CS-event was 7.5 minutes. In this situation, we observe that the light event effectively competes with the CS+/CS- association resulting in a selective and significant decrease in the freezing response to the CS- (Figure 3A/B). This is also appreciated in the correlation analysis (Figure 3C). The light event disrupts the correlation between the CS+ and CS-freezing responses, again showing that the association is effectively disrupted. As shown before, we do not observe any significant freezing response to light in the test trial, even if the light event is present temporally close to the CS+ even, suggesting that in our experimental conditions, memory association is not observed if events contain multimodal stimuli (Figure 3D).

**Figure 3.**
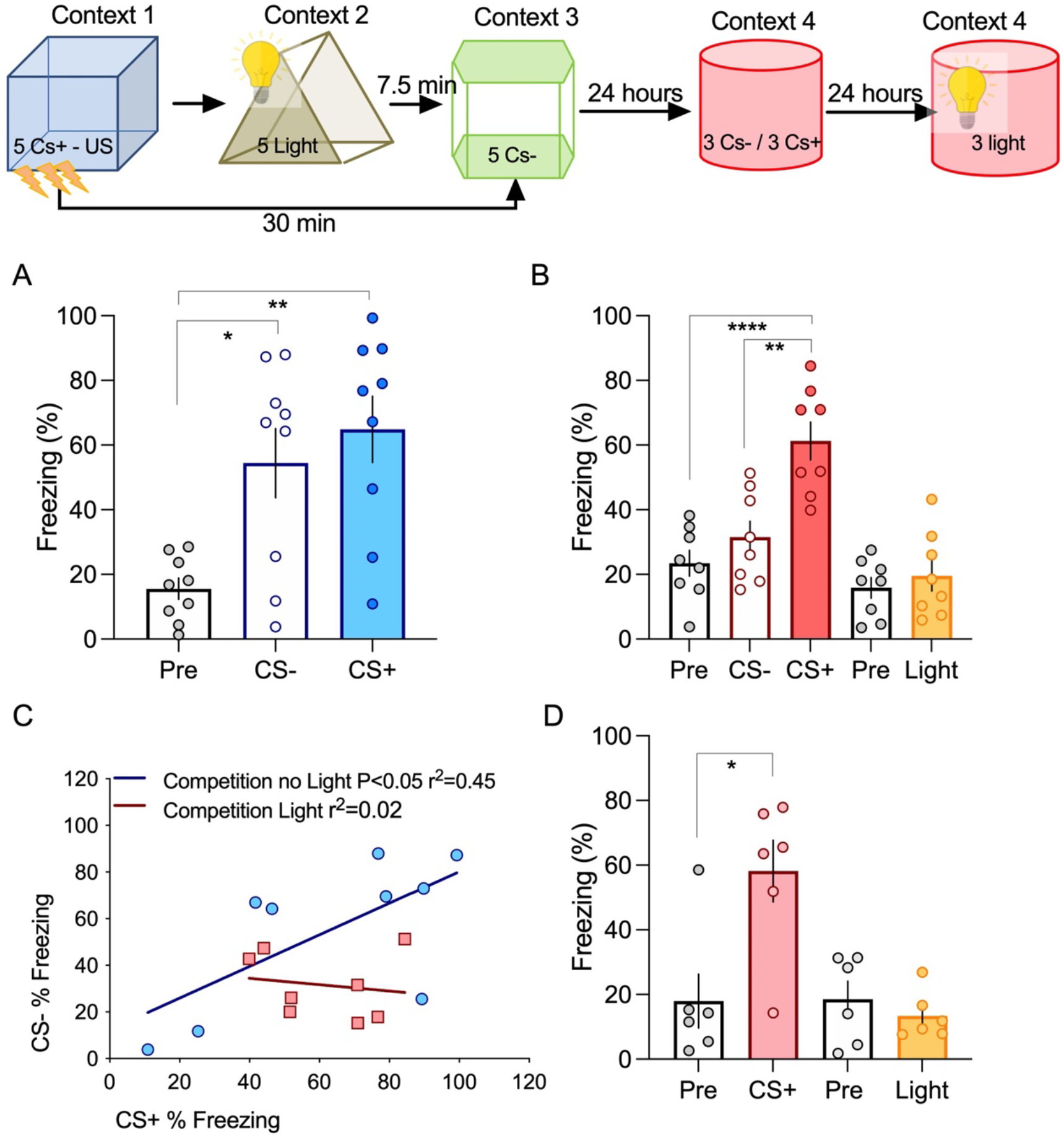
Exposing animals to a third event between event 1 and 2 leads to a stronger form of competition. A. Events are associated by cooperation if presented within a 30 minutes interval (ANOVA F=8.793 P=0.0014; *P=0.012; **P=0.0016 n=9). B. If animals are exposed to a third event, consisting of a different context paired with a light stimulus, between event 1 and 2, responses to the CS-significantly decrease (ANOVA F=16.02 P<0.0001; ****P<0.0001 **P=0.0011 n=8). C. The light event can effectively break the link between event 1 and 2, disrupting the correlation between fear responses to the CS+ and CS-. D. No association was seen between the fearful event and the light event, even if the CS-event is not present (Unpaired two-tailed t-Test pre-freezing vs CS+ *p=0.01; Unpaired two-tailed t-Test pre-freezing vs light p=0.43 n=6).

Given the multimodal properties of MGm neurons, we hypothesized that the light stimuli, which induces competition, can be mimicked by optogenetic stimulation of MGm axonal projections. To test this, we expressed a channelrhodopsin in the MGm, which resulted in strong labeling of axons in the internal commissure and the lateral amygdala (Figure 4A/A’). Light stimulation of the amygdala in the time interval between the CS+ and CS-, in control conditions (open vector) resulted in memory association and significantly higher freezing levels for both stimuli, as seen before (Figure 4B). Conversely, light stimulation in animals expressing channel rhodopsin in the thalamic inputs, led to competition, as seen by a significant decrease in the freezing response to the CS- (Figure 4C). This is also seen in the analysis of the discriminative index, which is higher in channel rhodopsin-expressing animals (Figure 5D). Interestingly, we found that light activation of thalamic inputs is sufficient to disrupt the correlation observed in control conditions, suggesting that activation of thalamic inputs efficiently modulates fear memory cooperation and competition (Figure 4E).

**Figure 4.**
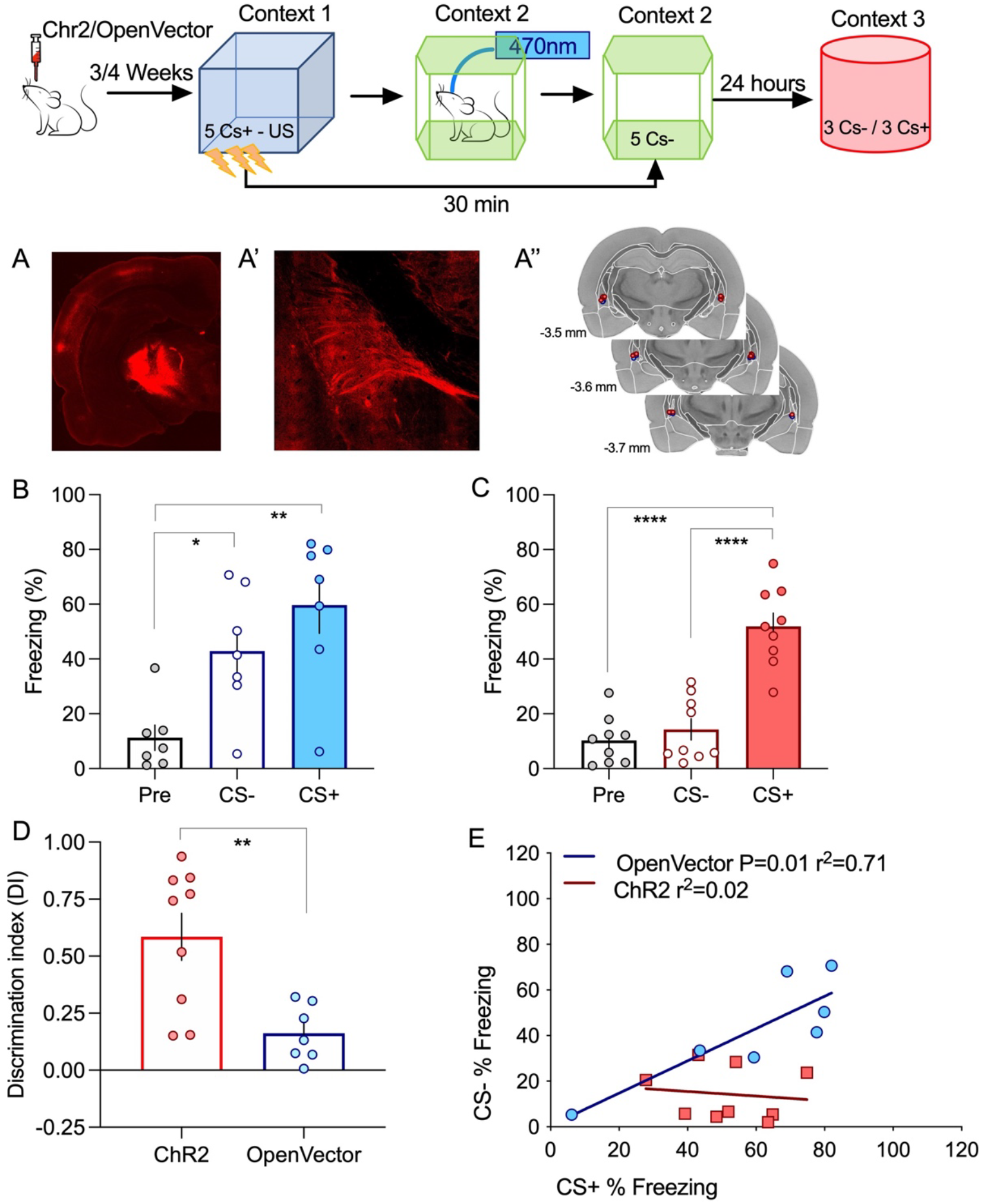
Memory competition can be induced by direct activation of the thalamic input A/A’. Injection of AAVs in the thalamic medial division of the medial genicular nucleus (MGm) results in transfection of afferents that project to cortical areas and the lateral amygdala, by the internal capsule. A’’ Location of the optical cannula used to deliver the light stimulation to thalamic inputs terminating in the lateral amygdala. B. Using the 30 minutes interval, light-activation of thalamic transfected with an open vector does not alter the ability of animals to associate between the neutral and aversive event (ANOVA F=9.001 P=0.002; *P=0.0352 **P=0.0016 n=7). C. Increasing thalamic activity by light-stimulation of ChR2 transfected inputs significantly decrease the response to the CS- (ANOVA F=34.17 P<0.0001; ****P<0.0001 ****P<0.0001 n=9). D. Light stimulation significantly increased the discrimination index (Unpaired two-tailed t-Test **p=0.0044). E. By inducing competition, light-stimulation of thalamic inputs effectively dissociates events 1 and 2 abolishing the correlation between CS+ and CS-responses observed in control conditions (open vector).

**Figure 5.**
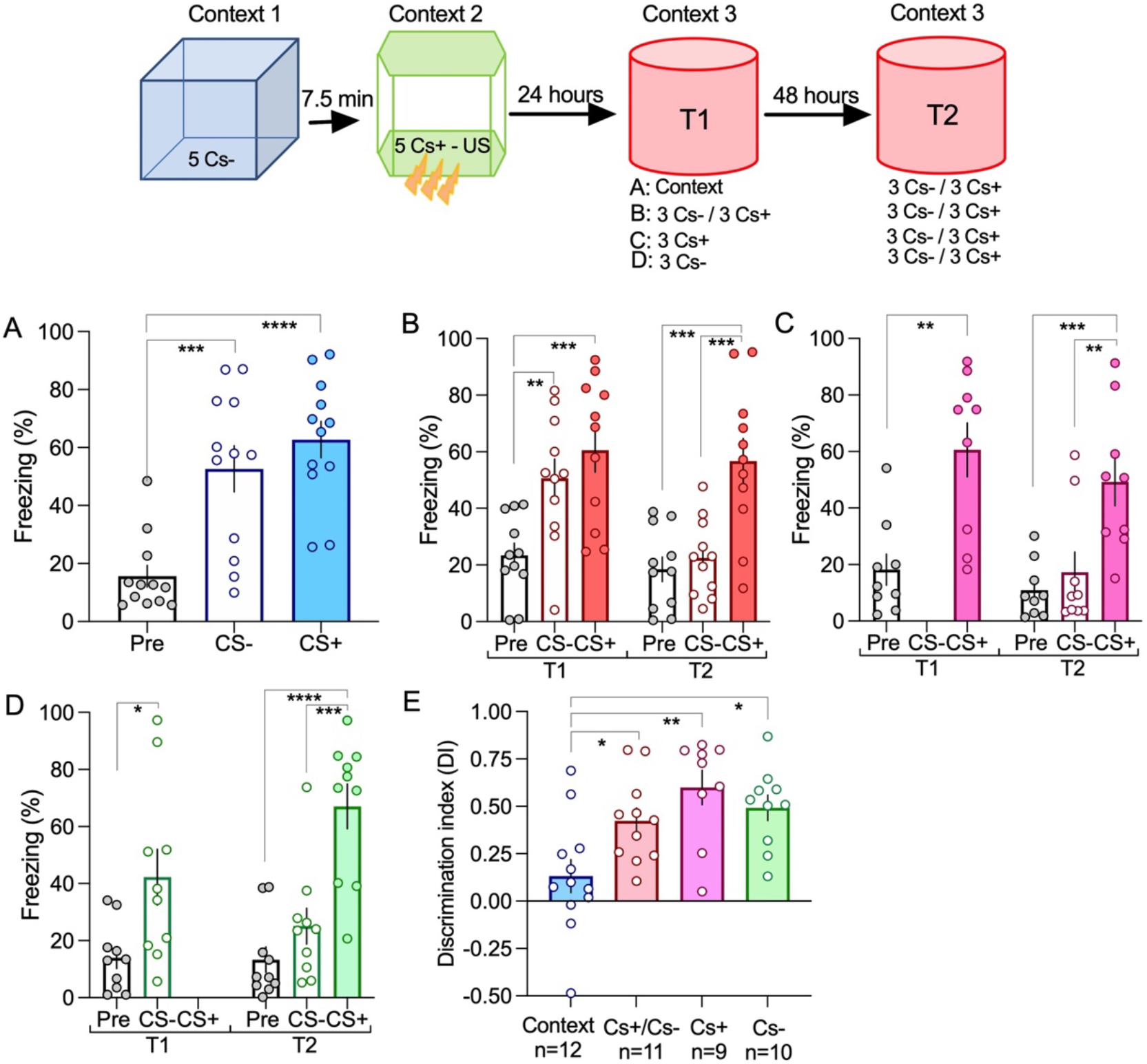
Memory reactivation leads to a selective decrease in the response to the neutral CS-event. A. Exposing animals to a new context does not induce reactivation and animals respond to the CS+ and CS-in subsequent test event (ANOVA F=15.86 P<0.0001; ***P=0.005 ****P<0.0001 n=12). B. Presenting the CS- and CS+ in the first test event leads to a significant reduction to the CS-in test 2 (Two-way repeated measures ANOVA F=7.69 P=0.0084; Time1 **P=0.0072 ***P=0.0002 Time2 ***P=0.0001 ***P=0.0006 n=11). C/D. Presenting the CS+ alone (C; Time1 Unpaired two-tailed t-Test **p=0.0014; Time2 ANOVA F=10.06 P=0.0007 ***P=0.001 **P=0.0047 n=9) or the CS-alone (D; Time1 Unpaired two-tailed t-Test *p=0.01; Time2 ANOVA F=19.73 P<0.0001 ****P<0.0001 ***P=0.0002 n=10) also resulted in a significant reduction to the CS-in test 2. E. Memory reactivation significantly increase the discrimination between fearful and neutral events (ANOVA F=6.539 P=0.0011 *P=0.047 **P=0.011 *P=0.012).

### Memory reactivation leads to the loss of the CS-freezing response

One of the essential features of memory is its ability to update. In our cooperation design, the memory trace represents the association between the CS+ and CS-events. Given that memory update requires reactivation of the previous trace, we designed a reactivation experiment in which the previous memory is either updated by the presentation of both sound stimuli (CS+ and CS-), only one of the sounds (CS+ or CS-) and no sounds (exposure to context only). Animals are then tested 48 hours after this reactivation trial, by exposing animal to the CS+ and CS-(test trial). We found that if animals are only exposed to context, no change is observed at the test trial, showing that exposure to context is not able to disrupt the previously associated memory (Figure 5A). In contrast, exposing animals to the auditory stimuli led to a significant decrease in the freezing response to the CS-, but not to the CS+ (Figure 5B/C/D). This reduction in the response to the CS-in the test trial is observed irrespectively of whether animals are exposed to both CS+ and CS- or only to the CS- or to the CS+. This is also appreciated by analyzing the discrimination index in the test trial. Only the condition where animals were only exposed to the context showed a significantly lower discrimination reflecting a high freezing response to both stimuli, CS+ and CS- (Figure 5E). These results show that reactivation is sufficient to trigger memory updating leading to a decrease of the non-significant stimuli response.

Fear memories are generally enduring^26^. To test whether our cooperative memory fades with time, we tested whether testing animals one month after training translates into a decrease in the freezing response to the auditory stimuli. We observe that one month after training, animals maintain their response to both the CS+ and CS- (Supplementary Figure 4A T1). As seen in the previous experiment, exposing animals to the auditory stimuli in the test trial led to a significant decrease in the CS-but not in the CS+ (Supplementary Figure 4A T2). This is also evident in the significant increase in the discrimination index in T2 as compared to T1 (Supplementary Figure 4B). This result shows that cooperative memories are enduring but very sensitive to disruption upon a single reactivation event.

## Discussion

The ability to link events, occurring in close temporal proximity, is highly relevant to acquiring associative memories. Equally important, or even more relevant, is the ability to separate events. Here, we addressed the temporal rules by which memories are associated or disconnected. Using a modified discriminative fear learning paradigm, we found that events occurring within short intervals, lower than 30 minutes result in the acquisition of an associative memory. If conversely, events occur separated by one hour they are not linked and thus, animals were able to discriminate between the fearful and the neutral stimuli, responding specifically to the fearful stimuli. Interestingly for the 30-minute interval, we found that animals were distributed in two distinct groups, animals that generalize by fearing both aversive and neutral stimuli and animals that do not respond to either. This observation suggests that for this intermediate time interval, in some animals, the neutral event can interfere with the fear memory acquisition of the aversive event. These temporal rules mirror the heterosynaptic interactions between thalamic and cortical inputs into the lateral amygdala, as described previously^24,25^. Interestingly, as stated in the introduction, we found that thalamic synapses are only able to cooperate with the cortical synapses if activated within 7.5 minutes^25^. This time restriction of thalamic cooperation is due to the activation of cannabinoid receptors (CB1R)^24,25^. Consistently, we also found that memory cooperation can be extended to 1 hour if CB1Rs are inhibited in the amygdala. Although we have previously shown that presynaptic CB1Rs are expressed in the thalamic pre-synaptic terminals and that CB1R agonists have a profound effect on thalamic excitatory synaptic transmission^24^, we cannot rule out a more generalized effect of CB1R antagonists, including modulation of inhibitory neurons and astrocytes that lead to an increase in arousal and thus fear expression^8,27,28^. We showed that increasing the activation of the thalamic input, by expressing an activating DREADD receptor increased memory cooperation leading to fear generalization. A similar manipulation of the cortical input did not alter the response of animals, which indicates that the increase in memory association is not due to a generalized increase in amygdala activation. It is noteworthy to further discuss that increasing the thalamic input activity not only increases memory association but it leads to a response similar to the cooperation 30 minutes, in which the response to the CS-is tightly correlated to the response to the CS+. Given that the correlation emerges because there are animals in which the CS-interferes with the fear memory of the CS-, this observation suggests that increasing thalamic activation increases the time window during which events interact but does not determine the direction of such interaction. These results go in line with a previous study that showed a critical role of post-learning activity of the thalamic input in the consolidation of fear memories^30^ and supports our hypothesis that thalamic activation determines whether animals associate or discriminate events.

Previous studies have shown that events can be associated in time^5,6^. Both these studies show that within a very large time window of 5 to 6 hours events can be associated and evoke a fearful memory, contrasting with the time rules that we report. In the first study, by Cai et al, the authors use a form of context fear conditioning, in which contexts are associated before the pairing of one of these contexts with a foot shock^5^. Context fear conditioning is dependent on the recruitment of the hippocampal network^9^ and thus different temporal rules may determine the linkage of hippocampal-dependent memories. Indeed, synaptic cooperation in hippocampal slices can occur in larger time intervals than the ones we described for the amygdala circuit^31,32^. In the second study, using auditory fear conditioning, the time interval is consistent with ours, given that in the case where the animals are not conditioned in the first event (CS alone) no cooperative effect was observed. For this condition, the shortest interval tested by the authors was 90 minutes, but neither interval renders a positive cooperative effect on the second event^6^. Additionally, we have found that heterosynaptic cooperation between thalamic and cortical synapses follows different temporal rules in rats and mice ^29^, which can translate in different time rules for memory association.

Our temporal rule also applies to memory competition. We found that exposing animals to a competitive event, after auditory fear memory association only destabilizes previous associative memory if occurring within a short time interval. The competitive effect is mild and it is important to note that in our experiment setting the light neutral event was never associated with the aversive event. These results suggest that in our experimental design, we do not find a multimodal memory association (CS+ - light association) which might explain the mild competitive effect observed. Importantly, when the competitive event occurs between the two auditory events, we observe that memory linkage is abolished, shown by eliminating the correlation between the CS- and CS+ response. As for cooperation, optogenetic activation of the thalamic input is sufficient to mimic the light competition, disrupting the association between the neutral and the fearful event. The effect of activating thalamic input is different whether one is in a cooperative or competitive scenario. For the 30-minute interval, we see an overall cooperative effect although, as stated above, some animals show competition. In this scenario, further activation of the thalamic input favors competition increasing discrimination. For the 60-minute interval, in control conditions, there is no cooperation and thus, activation of the thalamic input increases cooperation.

Although enduring, the fear-associative memory for the neutral event is very sensitive to reactivation. Re-exposing animals to any of the auditory stimuli, leads to the loss of the response to the CS-, showing that reactivation promotes fear discrimination. Our results resonate with a previous study showing that a distractor at the time of memory reactivation can induce a competitive disruption of a previously acquired fear memory^33^. This suggests that once reactivated, memory is updated towards maintaining the most relevant event. Taken together, we show a clear time rule determining whether events are associated by cooperation or interfere by competition in fear memory acquisition and maintenance. We also show that the modulation of thalamic synaptic activity has a critical role in determining this temporal rule. Given the individual differences in memory association and the proposed role of the endocannabinoid signaling system in its modulation, we propose that individual differences in the endocannabinoid system may explain individual susceptibilities to stress.

## Acknowledgments

We would like to thank Ana Carolina Temporão for developing the scripts used to run the behavioral experiments. We would also like to acknowledge the support of the Imaging facility at i3S and the Animal Facility both at i3S and NOVA Medical School. This study was supported by a Fundação para a Ciência e Tecnologia (FCT) Project Grant (PTDC/MED NEU/30772/2017 LISBOA-01-0145-FEDER-030772) and a Grant from the Brain and Behaviour Foundation to RF (NARSADYI-2016-GA25118). RF was supported by an CEEC individual grant (2020.02221.CEECIND; DOI 10.54499/2020.02221.CEECIND/CP1586/CT0013), NM was supported by a FCT PhD fellowship (SFRH/BD/130911) and IC is supported by research grants in the scope of the Fundação para a Ciência e Tecnologia (FCT) Project Grant (2022.09002.PTDC, DOI 10.54499/2022.09002.PTDC).

## Contributions

Conceptualization: RF; Data Curation: RF, NM, IC; Data Analysis: RF, NM, IC; Funding acquisition and Project administration: RF; Supervision: RF; Writing – original draft: RF, NM; Writing – review & editing: RF

## Supplementary Material

### Materials and Methods

#### Animals

Naïve male Sprague-Dawley rats (200-350gr) obtained from a commercial supplier (Charles River Laboratories, France) or an in-house breeding stock at NOVA Medical School animal facility, were used in this study. Animals were allowed to acclimate after transportation for at least one week after arrival. Rats were housed in a temperature- and humidity-controlled housing facility under a 12h light-dark cycle (lights ON at 8 a.m. OFF at 7 p.m.), with ad libidum access to food and water. All the behavioral procedures were performed during the light phase of the cycle. Animals were handled for two days before each experiment. Rats were housed two per cage except between training and the first testing session in which animals were housed alone. In the experiments where testing was performed with 30 days of interval from training, animals were housed alone for 24 hours (after training) and then in pairs for the remaining duration of the behavioral trial. All procedures that involved the use of rats were approved by the Portuguese Veterinary Office (Direcção Geral de Veterinária e Alimentação — DGAV) and the ethics committee of NOVA Medical School.

#### Fear conditioning paradigm

All the behavioral studies were performed in sound-attenuating boxes present in the same procedure room. A total of four different conditioning boxes were used for conditioning and testing sessions. To minimize generalization between contexts, the chambers differ in size, shape, color, odor, lighting conditions, and floor composition. In context A the walls are made of black acrylic plates and were ornamented with yellow triangles. Before and after each experiment, the chamber was cleaned with 1% lavender detergent. In context B, the walls were made of clear Plexiglass, were ornamented with yellow circles, and were cleaned with 1% acetic acid. In context C, the walls are made of black acrylic plates, ornamented with white stripes, and cleaned with 1% marine detergent. In context D, the walls were made of clear Plexiglass, ornamented with black stripes, and cleaned with 70% ethanol. All the chambers contained an electrical grid floor (that is covered during the CS-presentation and probe test); a speaker through which auditory stimuli were delivered, a house light, a fan, and infra-red light. The different chambers were used in a counterbalanced manner across experiments.

##### Memory cooperation experiments

Animals were exposed to two tones, one paired with a foot-shock (CS+: continuous auditory tone at 2 kHz, 10-ms rise and fall, 75dB, 20 sec) and one unpaired (CS-: auditory pips, 10 kHz pips repeated at 0.5Hz, 10-ms rise and fall, 75dB, 20 sec). Rats were placed in a conditioning chamber and allowed to explore for 2 minutes before any auditory tone was presented. After this, animals were exposed to a block of five presentations of the CS- (in a pseudo-random manner, iTi 60-180s). After different time intervals – 7.5 minutes, 30 minutes, or 1 hour – animals were transferred to another context and conditioned with a block of five presentations of CS+ that co-terminated with a foot shock (1mA, 1s). After training, rats remained in the chamber for an additional 30s before returning to their home cage. On day 2, animals were tested (probe test) in a novel chamber by presenting three trials of the CS-followed by three trials of the CS+ (same settings as in training, pseudo-random presentation, iTi 60-180s). In the case of the reactivation and remote experiments, animals were tested twice, on consecutive days. In that case, the same context was used for testing (T1 and T2).

##### Memory competition experiments

In memory competition experiments, we used a light stimulus (Green led, Intensity: 35 LUX, Duration: 0.48sec; Interval: 0.86sec) as a CS3. As before, we used two auditory stimuli, CS- and CS+ with the same parameters as in cooperation experiments. In this experimental setting, the CS+ was presented first, after which the animal transitioned to another context where it remained until the second event initiated (five presentations of the CS-; see diagram in figures). The light stimulus was presented in a third context either after the CS-presentation (first experiment) or in between the CS+ and CS-events. The total time interval between CS+ and CS-events was always 30 minutes. On day 2, animals were tested for CS- and CS+ stimuli, as described for cooperation experiments. The light probe test occurred on day 3, by presenting three trials of the light stimuli in the same context as in probe test 1.

#### Viruses

Adeno-associated viral vectors (AAV) were purchased from Addgene. For the optogenetic experiments, we used serotype 1, pAAV-CaMKIIa-hChR2(H134R)-mCherry, a gift from Karl Deisseroth (Addgene plasmid#26975; http://n2t.net/addgene:26975;RRID:Addgene_26975), titer 1×10^13^ vg/mL and the serotype 1, pAAV-CaMKIIa-mCherry a gift from Karl Deisseroth (Addgene plasmid#114469; http://n2t.net/addgene:114469;RRID:Addgene_114469), titer 1×10^13^ vg/mL. For the chemogenetic experiments we used the serotype 9, pAAV-CaMKIIa-hM3D(Gq)-mCherry a gift from Bryan Roth (Addgene plasmid # 50476;http://n2t.net/addgene:50476;RRID:Addgene_50476), titer 1×10^13^ vg/mL and serotype 9, pAAV-CaMKIIa-mCherry a gift from Karl Deisseroth (Addgene plasmid#114469; http://n2t.net/addgene:114469;RRID:Addgene_114469), titer 1×10^13^ vg/mL.

#### Surgery and Viral transfection

Rats, 6-8 weeks of age were anesthetized with isoflurane (4%; (SomnoSuite anesthesia system, Kent Scientific) and maintained at 1.5-2% throughout the surgery in the stereotaxic setup (Kopf 940, Kopf Instruments). Surgery started with a skin incision, after local injection of 2% lidocaine, and retraction to expose the skull, where a craniotomy and durotomy were performed. AAVs were delivered bilaterally into the target brain areas using a pulled glass pipet (tip diameter 20-30um) connected to a manual injector (WPI instruments). A volume of 0.3uL (for A1) or 1uL (for MGm), per hemisphere was injected, over 10 minutes. After injecting the virus, the needle was left at the site of injection for an additional 10 minutes to allow the virus to diffuse into the tissue. After that, the skin was sutured. Coordinates (from bregma) for targeting the different brain areas were the following: auditory cortex (A1): -3.6 (AP); ± 6.4 (ML); -5.0 (DV from the skull surface); MGm: -5.28 (AP); ± 3.40 (ML); -6.20 (DV from the skull surface), LA: -3.00 (AP); ± 5.20 (ML); -7.20 (DV from the skull surface); all in mm, according to Paxinos and Watson (1986). In the case where implants were needed (optogenetics and in vivo pharmacology), the implants were fixed to the skull with anchored screws, medical glue (Vetbond), and dental cement (Dentalon Plus, Kulzer). Optic fiber cannulas were lowered to 7.2DV below the skull surface. Thirty minutes before the end of the surgery, animals were administered carprofen (5mg/Kg) and 1mL of saline solution for hydration. Animals were then placed in a heated cage post-surgery for recovery. For the first, 2 days postoperative, carprofen was administered. For optogenetic and chemogenetic manipulations, rats had 4-5 weeks of recovery to allow adequate virus expression.

#### In-vivo pharmacology

For in vivo pharmacology, the experiments were done 1 week after the guide cannula implantation. Guide cannulas were implanted over the amygdala, using the same AP and ML coordinates as described previously. Rats were implanted bilaterally with a 7.2mm guide cannula (26 gauge; PlasticsOne, Bilaney). The guide cannula was fixed to the skull using dental acrylic and two bone screws. Dorsoventral coordinates were measured from the skull surface with the internal cannula (33 gauge; PlasticsOne, Bilaney) extending 0.5mm beyond the end of the guide cannula. Intracranial infusion was administered using an injection needle (Plastic One, USA) inserted through the guide cannula with the injection needle connected to 0.5-μL syringes with polyethylene tubes and controlled by an automated microinjection pump (Precision Pump, Longer). AM281 solution (25μM, HelloBio), diluted in 1% of DMSO, of a volume of 200nL was injected 5min after the end of CS^-^ presentation at a rate of 20nL/min. The control animals were infused with only vehicle solutions.

#### Chemogenetics

Adeno-associated viral vectors expressing the hM3D(Gq) receptor (described above) were injected either in the MGm thalamic region or in the primary auditory cortex, both areas projecting to the LA region of the amygdala. A 3-4 weeks expression time was given to allow a stable expression in the axons. After this period, for three days, rats were habituated to intraperitoneal (IP) injections of saline before CNO (4mg/Kg, HelloBio) was administered, 40 minutes before the training session. After each experiment, the verification of the transfection area was performed by PFA brain fixation and cryostat brain slicing.

#### Optogenetics

Channel rhodopsin expressing AAV and open vectors (described above) were injected in the MGm thalamic region projecting to the LA, and a 3-4 weeks expression time was given to allow a stable expression in the axons. Two 200-micrometer (Thorlabs Fiber Optic Cannula, 2.5mm OD Ceramic, 200um, 0.39NA, L=10mm) optic fibers were implanted in the same surgery to target LA. Fibers were inserted and glued to a cannula guide (Thorlabs OGF-5) and lowered to 7.1mm below the skull surface. To stimulate ChR2, a 470 nm high-power LED light source was used with a light intensity of 7mW/mm^2^. The end of each optical fiber was connected to the LED light source (470 nm, BLS-LCS-0470 Mightex Systems, Toronto, Canada), controlled by an LED driver (BLS-SA04-US Mightex Systems) and a computer. Transfected terminals in the amygdala are light-stimulated with 30 pulses at 0.033 Hz, 75% of LED intensity, delivered bilaterally, during the interval between the CS- and CS+. All the freely moving experiments were done with a rotary joint (Thorlabs). After each experiment, the verification of the optic fiber tract and area of transfection was performed by PFA brain fixation and cryostat brain slicing.

#### Imaging

At the end of the experiments, animals received an overdose of isoflurane and the brains were removed from the skull and fixed in buffered 4% paraformaldehyde (pH=7.5), overnight. Brains were then sectioned with a cryostat (Leica CM3050 S) in sections of 40μm thickness. Imaging was performed using a BC 43 bench confocal microscope (Andor, Oxford Instruments). Whole brain slides tile images were taken using a 2.5X (0,06) or 20X objective (0,8). The emission wavelength for mcherry was 590 nm with 650 ms of exposure time. Brain slices were inspected to confirm the virus expression in the thalamic and cortical regions projecting to the LA, as well as expression within amygdala areas. Brightfield images were taken to determine the cannula or optic fiber location in the LA.

### Data collection and Statistical analysis

Animal behavior was recorded using a video camera. A semi-automated video tracking software (Bonsai®) was used to quantify behavioral scores. Behavioral performance was measured based on the percentage of freezing to baseline responses (pre-freezing). Freezing was scored from the video recording using a frame-by-frame analysis of pixel changes. Freezing responses evoked by the three auditory stimuli were averaged. Pre-freezing corresponds to the average freezing of the 120 seconds before tone presentation. Animals that showed a pre-freezing response above the mean+SD within the group were excluded. Fear discrimination index (DI) was calculated by 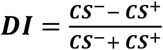. Statistical analyses were done via Prism (GraphPad Software, San Diego, CA, USA). All the data are represented as mean ± SEM. Before choosing the statistical test, a normality test (Shapiro-Wilk normality test) was done on all data sets. If the data presented a normal distribution, then a parametric test (one-way ANOVA with Tukey’s post hoc test for multiple comparisons) was used to calculate the statistical differences between groups. In the case of the reactivation and remote experiments, given that animals are tested two times, a repeated-measures ANOVA was used with post-hoc correction of significant interactions. The main effects and P values are mentioned in the figure legends. For the correlation analysis, we plotted the percentage freezing to the CS+ versus the percentage freezing to the CS- and performed a Pearson correlation analysis. R squares and p-values are depicted in figures.

## Supplementary Figures

**Supplementary Figure 1.**
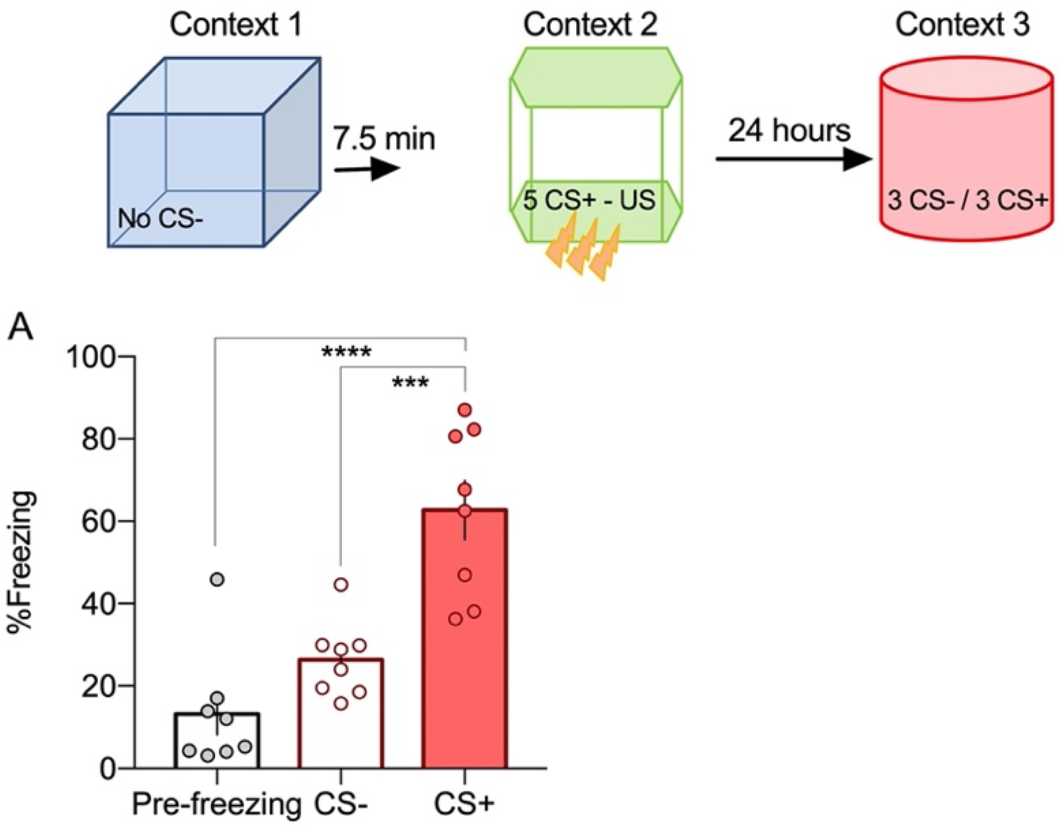
A. Memory association is not observed if the CS-auditory stimulus is not present during the training (ANOVA F=22.64 P<0.0001; ****P<0.0001 ***P=0.0003 n=8).

**Supplementary Figure 2.**
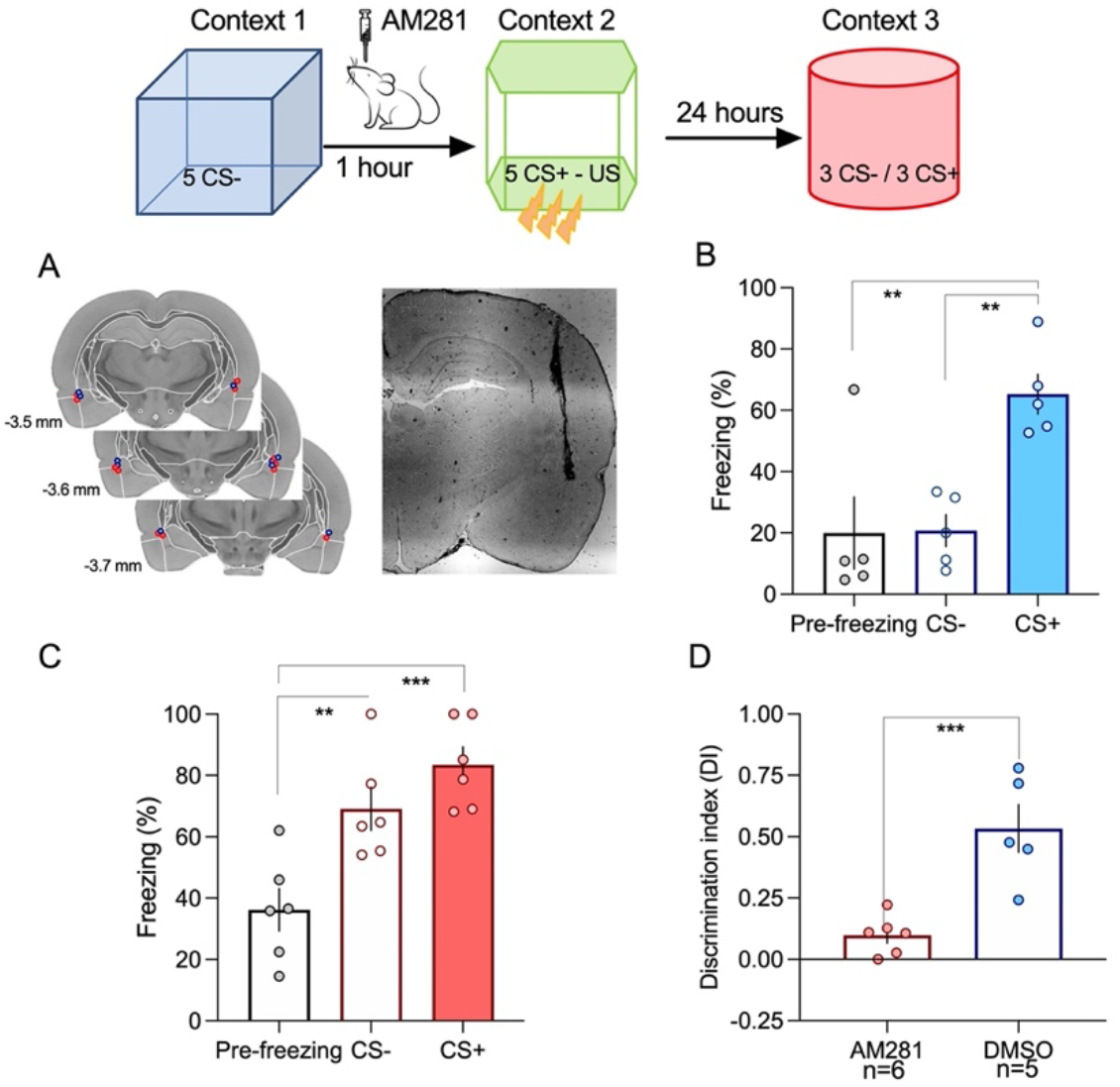
Inhibiting CB1 receptors by AM281 application in the lateral amygdala increases fear memory association. A. Placement of internal cannula. B. In control conditions, animals do not associate events separated by one hour (ANOVA F=9.7 P=0.0031; **P=0.006 **P=0.006 n=6). C. Inhibition of CB1R significantly increases fear responses for the neutral event (ANOVA F=13.47 P=0.0004; **P=0.008 ***P=0.0004 n=5). D. Discrimination is significantly decrease if CB1R are inhibited (Unpaired two-tailed t-Test **p=0.0013).

**Supplementary Figure 3.**
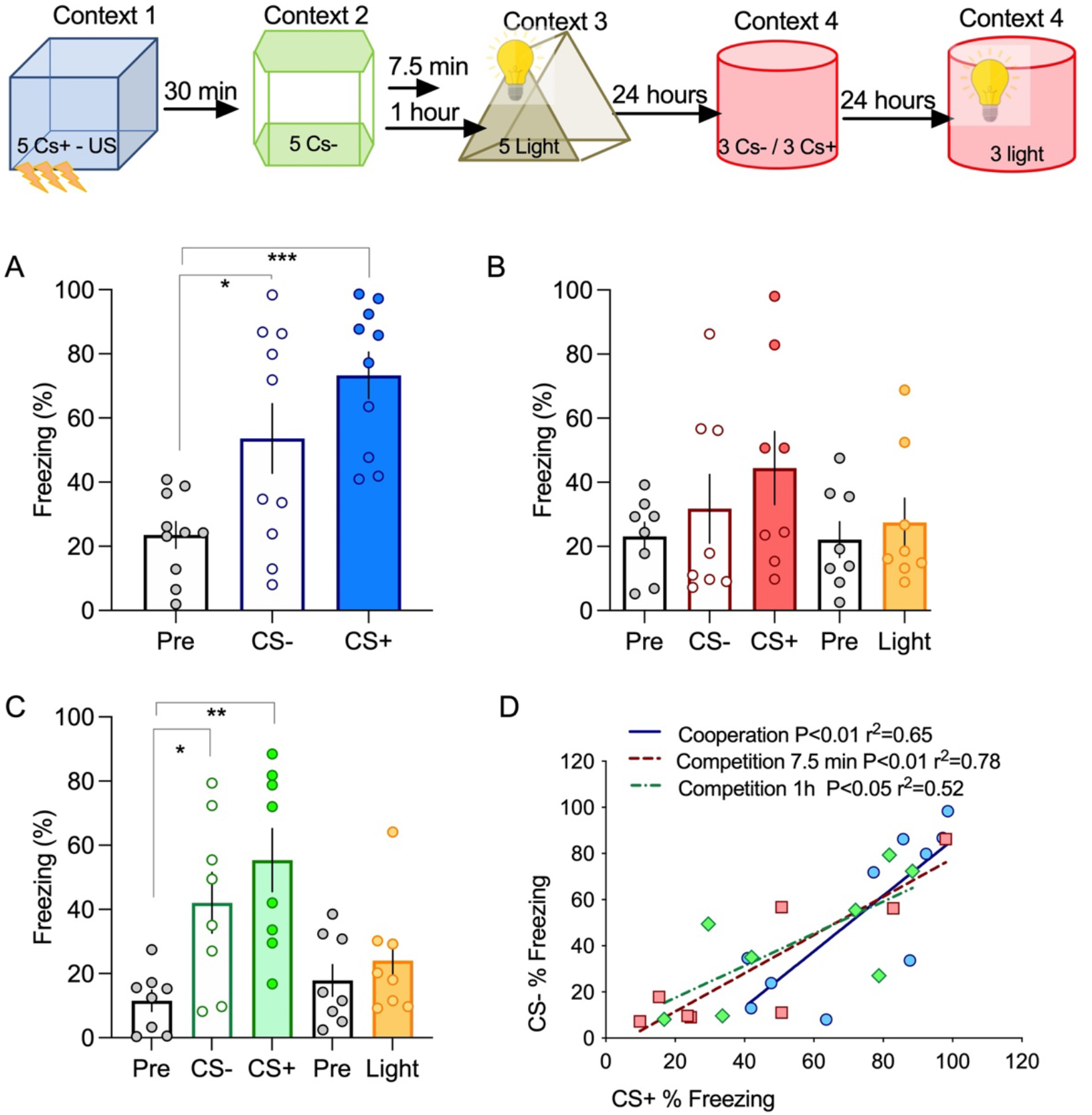
Exposing animals to a third event leads to competition. A. Events are associated by cooperation with a 30 minutes interval (ANOVA F=10.04 P=0.0005; *P=0.031 **P=0.0004 n=10). B. If animals are exposed to a third event, consisting of a different context paired with a light stimulus within 7.5 minutes, responses to the CS+ and CS-decrease and are no longer different from pre-freezing levels (ANOVA F=1.3 P=0.29 n=8). C. Increasing the time interval between event 2 and 3 to 60 minutes prevents competition (ANOVA F=7.67 P=0.0031; *P=0.037 **P=0.0027; Unpaired two-tailed t-Test pre-freezing vs Light p=0.46 n=8) D. Although the third event induces competition, the responses to the CS+ and CS-are still linked. All conditions show a positive correlation indicating the strong link between event 1 and 2.

**Supplementary Figure 4.**
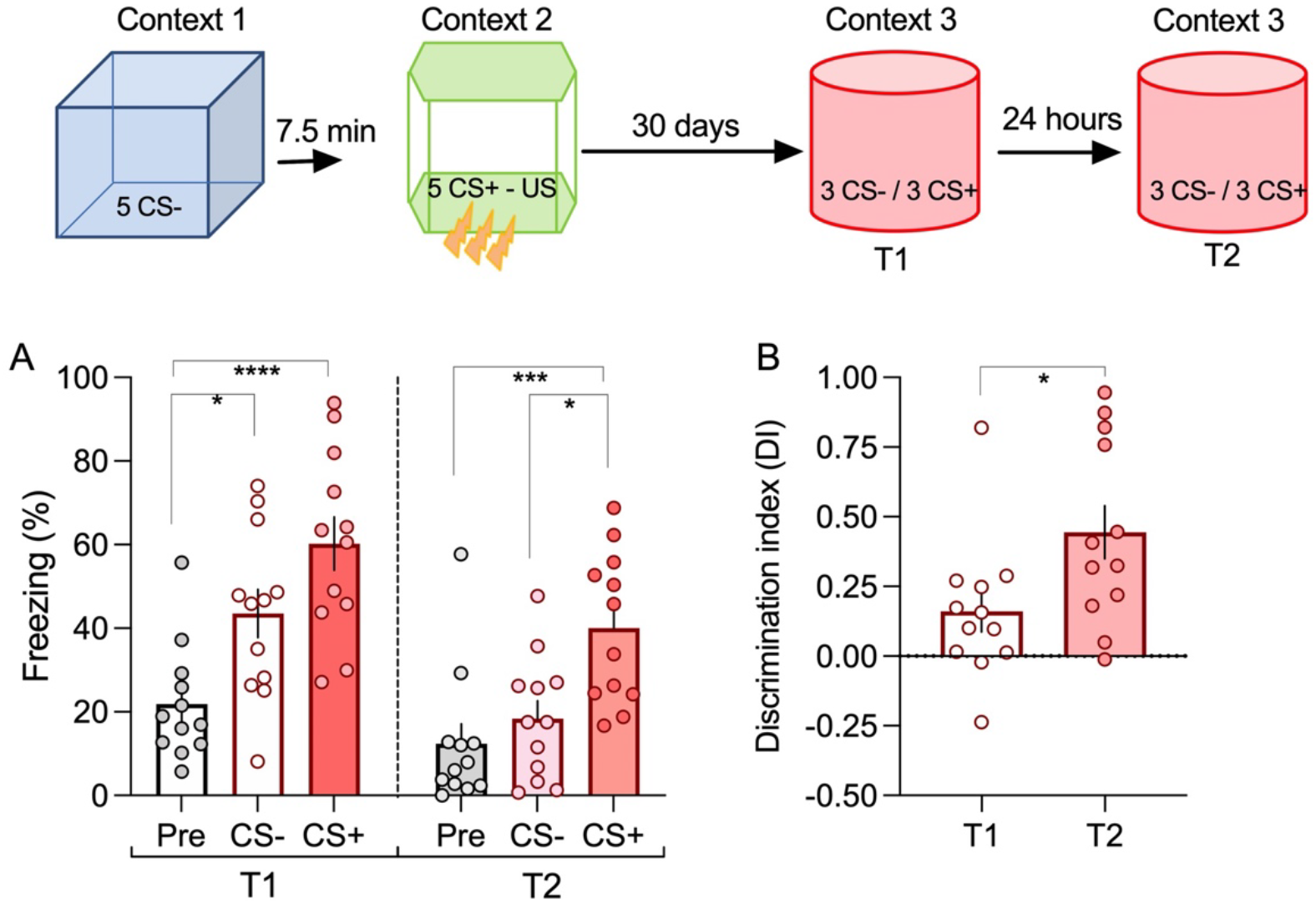
Memory association is enduring. A. Animals show an intact memory association even if tested one month after training. As before, reactivation of memory led to a significant decrease in the response to the neutral (CS-) event (Two-way repeated measures ANOVA F=51.36 P<0.0001; Time 1 *P=0.01 ****P<0.0001 Time 2 ****P<0.0001 ***P=0.0002 n=12). B. Memory reactivation significantly increased discrimination between the fearful and neutral event, by selectively decrease the fearful response to the neutral event (Unpaired two-tailed t-Test *p=0.027).

## References

1. McKenzie, S. & Eichenbaum, H. Consolidation and reconsolidation: two lives of memories? Neuron 71, 224–33 (2011).

2. Lee, J. L. C. C. Reconsolidation:maintaining memory relevance. Trends Neurosci. 32, 413–420 (2009).

3. Pedreira, M. E., Pérez-Cuesta, L. M. & Maldonado, H. Mismatch Between What Is Expected and What Actually Occurs Triggers Memory Reconsolidation or Extinction. Learn. Mem. 11, 579–585 (2004).

4. Forcato, C., Rodríguez, M. L. C., Pedreira, M. E. & Maldonado, H. Reconsolidation in humans opens up declarative memory to the entrance of new information. Neurobiol. Learn. Mem. 93, 77–84 (2010).

5. Cai, D. J. et al. A shared neural ensemble links distinct contextual memories encoded close in time. Nature 534, 115–118 (2016).

6. Rashid, A. J. et al. Competition between engrams influences fear memory formation and recall. Science (80-.). 353, 383–387 (2016).

7. Moncada, D. & Viola, H. Induction of long-term memory by exposure to novelty requires protein synthesis: evidence for a behavioral tagging. J. Neurosci. 27, 7476–81 (2007).

8. Fonseca, R., Madeira, N. & Simoes, C. Resilience to fear: The role of individual factors in amygdala response to stressors. Molecular and Cellular Neuroscience (2021). doi:10.1016/j.mcn.2020.103582

9. Johansen, J. P., Cain, C. K., Ostroff, L. E. & Ledoux, J. E. Molecular mechanisms of fear learning and memory. Cell (2011). doi:10.1016/j.cell.2011.10.009

10. Kim, J. J. & Jung, M. W. Neural circuits and mechanisms involved in Pavlovian fear conditioning: A critical review. Neuroscience and Biobehavioral Reviews (2006). doi:10.1016/j.neubiorev.2005.06.005

11. Ghosh, S. & Chattarji, S. Neuronal encoding of the switch from specific to generalized fear. Nat. Neurosci. 18, 112–20 (2015).

12. Blair, H. T., Schafe, G. E., Bauer, E. P., Rodrigues, S. M. & LeDoux, J. E. Synaptic plasticity in the lateral amygdala: a cellular hypothesis of fear conditioning. Learn. Mem. 8, 229–242 (2001).

13. Bordi, F. & LeDoux, J. E. Response properties of single units in areas of rat auditory thalamus that project to the amygdala. I. Acoustic discharge patterns and frequency receptive fields. Exp. brain Res. 98, 261–74 (1994).

14. Kimura, A., Donishi, T., Sakoda, T., Hazama, M. & Tamai, Y. Auditory thalamic nuclei projections to the temporal cortex in the rat. Neuroscience 117, 1003–1016 (2003).

15. Han, J.-H. et al. Increasing CREB in the auditory thalamus enhances memory and generalization of auditory conditioned fear. Learn. Mem. 15, 443–453 (2008).

16. Antunes, R. & Moita, M. A. Discriminative auditory fear learning requires both tuned and nontuned auditory pathways to the amygdala. J. Neurosci. 30, 9782–7 (2010).

17. Pinho, J., Marcut, C. & Fonseca, R. Actin remodeling, the synaptic tag and the maintenance of synaptic plasticity. IUBMB Life (2020). doi:10.1002/iub.2261

18. Fonseca, R., Nägerl, U. V., Morris, R. G. M. & Bonhoeffer, T. Competing for memory: hippocampal LTP under regimes of reduced protein synthesis. Neuron 44, 1011–20 (2004).

19. Sajikumar, S., Morris, R. G. M. & Korte, M. Competition between recently potentiated synaptic inputs reveals a winner-take-all phase of synaptic tagging and capture. Proc. Natl. Acad. Sci. U. S. A. 111, 12217–21 (2014).

20. Szabó, E. C., Manguinhas, R. & Fonseca, R. The interplay between neuronal activity and actin dynamics mimic the setting of an LTD synaptic tag. Sci. Rep. 6, 33685 (2016).

21. Fonseca, R. Synaptic cooperation and competition: Two sides of the same coin? Synaptic Tagging and Capture: From Synapses to Behavior (2015). doi:10.1007/978-1-4939-1761-7_3

22. Drumond, A., Madeira, N. & Fonseca, R. Endocannabinoid signaling and memory dynamics: A synaptic perspective. Neurobiol. Learn. Mem. 138, 62–77 (2017).

23. Fonseca, R. Asymmetrical synaptic cooperation between cortical and thalamic inputs to the amygdale. Neuropsychopharmacology 38, 2675–87 (2013).

24. Madeira, N., Drumond, A. & Fonseca, R. Temporal Gating of Synaptic Competition in the Amygdala by Cannabinoid Receptor Activation. Cereb. Cortex (2020). doi:10.1093/cercor/bhaa026

25. Fonseca, R. Asymmetrical synaptic cooperation between cortical and thalamic inputs to the amygdale. Neuropsychopharmacology (2013). doi:10.1038/npp.2013.178

26. Schafe, G. E., Nader, K., Blair, H. T. & LeDoux, J. E. Memory consolidation of Pavlovian fear conditioning: A cellular and molecular perspective. Trends in Neurosciences (2001). doi:10.1016/S0166-2236(00)01969-X

27. Lutz, B., Marsicano, G., Maldonado, R. & Hillard, C. J. The endocannabinoid system in guarding against fear, anxiety and stress. Nat. Rev. Neurosci. 16, 705–718 (2015).

28. Jacob, W., Marsch, R., Marsicano, G., Lutz, B. & Wotjak, C. T. Cannabinoid CB1 receptor deficiency increases contextual fear memory under highly aversive conditions and long-term potentiation in vivo. Neurobiol. Learn. Mem. 98, 47–55 (2012).

29. Faress, I. et al. Non-Hebbian plasticity transforms transient experiences into lasting memories. (2023). doi:10.1101/2023.04.06.535862

30. Lee, Y., Oh, J. P. & Han, J. H. Dissociated role of thalamic and cortical input to the lateral amygdala for consolidation of long-term fear memory. J. Neurosci. (2021). doi:10.1523/JNEUROSCI.1167-21.2021

31. Redondo, R. L. & Morris, R. G. M. Making memories last: the synaptic tagging and capture hypothesis. Nat. Rev. Neurosci. 12, 17–30 (2011).

32. Redondo, R. L. et al. Synaptic tagging and capture: differential role of distinct calcium/calmodulin kinases in protein synthesis-dependent long-term potentiation. J. Neurosci. 30, 4981–9 (2010).

33. Crestani, A. P. et al. Memory reconsolidation may be disrupted by a distractor stimulus presented during reactivation. Sci. Rep. (2015). doi:10.1038/srep13633

